# Recapitulating whipworm development *in vitro* using caecaloids

**DOI:** 10.64898/2026.03.16.712207

**Authors:** Denise Tran, Charlotte Tolley, Thomas Morris, Emily Hart, Matthew Berriman, Stephen R. Doyle, Maria A. Duque-Correa

**Author notes:** Corresponding author (M. A. Duque-Correa). Magdalene College, University of Cambridge, CB3 0AG, United Kingdom (Thomas Morris). Anne McLaren Building, 90 Francis Crick Avenue, Cambridge Biomedical Campus, University of Cambridge, CB2 0BA, United Kingdom (Emily Hart). These authors contributed equally to this work (Denise Tran, Charlotte Tolley).

## Abstract

Whipworms (*Trichuris* spp.) are intracellular intestinal parasites that develop within the host caecal epithelium, yet the host signals that regulate their growth and developmental progression remain poorly understood. Progress in studying these processes has been limited by the lack of physiologically relevant *in vitro* systems capable of supporting sustained whipworm development. Here, we established an *in vitro* infection system using caecal organoids (caecaloids) and evaluated their capacity to support sustained growth and morphological development of *Trichuris muris* larvae. To rigorously validate this system, we generated a comprehensive and up-to-date anatomical and biometrical reference dataset describing the whole-body growth and tissue-level morphogenesis of *T. muris* throughout its life cycle *in vivo*. Quantitative analysis across larval and adult stages confirmed that the trajectory of parasite growth is largely conserved across host mouse strains and provided a detailed contextualised description of the development of key anatomical structures of *T. muris*. Using this reference framework, we evaluated parasite growth and development in long-term *T. muris*–caecaloid co-cultures. Larvae invading the caecaloid epithelium remained intracellular within syncytial tunnels and exhibited sustained growth over extended culture periods. *in vitro* parasites developed increasing anatomical complexity, including formation of the bacillary band, stichosome, intestine, and rectum. Importantly, quantitative comparisons revealed that larvae developing within caecaloids follow growth trajectories and morphological developmental patterns closely resembling those observed *in vivo*. This study therefore presents the first detailed anatomical and morphometric framework for validating whipworm development in an organoid system and provides concrete evidence that the caecaloid epithelium is sufficient to trigger and sustain whipworm growth and morphogenesis, establishing caecaloids as a powerful experimental platform for investigating *Trichuris* infection and development.

## Introduction

*Trichuris* is a speciose genus of parasitic roundworms with more than 70 species that infect humans and other mammals. Human-infective whipworms (*Trichuris trichiura*) are soil-transmitted helminths that infect hundreds of millions of people worldwide. Chronic *Trichuris* infections result in trichuriasis, which is a major cause of neglected tropical disease-associated morbidity, associated with malnutrition, growth stunting, anaemia, and impaired cognitive development, imposing substantial socio-economic burdens on affected countries (Jourdan et al., 2018; Else et al., 2020; Behniafar et al., 2024). Despite this global impact, no vaccines for whipworm are available, and current control strategies rely on chemotherapeutic interventions that often fail to clear infections (Moser et al., 2017; Welsche et al., 2023; Gebreyesus et al., 2024). A deeper understanding of whipworm biology is therefore urgently needed to inform the development of new therapeutic and preventive strategies against whipworm infections.

*Trichuris* spp. occupy a highly distinctive niche within the epithelium of the caecum and proximal colon of their hosts. Whipworm life cycle is direct and confined to a single host with no free-living stage. Infection begins when embryonated eggs are ingested and hatch in the caecum in a process mediated by the host microbiota (Klementowicz et al., 2012; Else et al., 2020; Mkandawire et al., 2022). Motile first-stage (L1) larvae transverse the mucus layers and penetrate the intestinal epithelial cells (IECs) at the bottom of the crypts of Lieberkühn, where they establish an intracellular syncytial niche and undergo successive moults through larval stages before reaching adulthood (Else et al., 2020; Duque-Correa et al., 2022). From the L3 stage onwards, parasites are located at the top of the crypts, with the anterior section remains embedded within IECs while their posterior end protruding into the gut lumen. This enables mating and the production of eggs, which are shed in host faeces to complete the life cycle (Klementowicz et al., 2012; Hurst and Else, 2013; Else et al., 2020).

*T. trichiura* is an obligate human parasite and cannot be maintained in the laboratory settings. As a result, much of our understanding of whipworm biology and host–parasite interactions has relied on animal models, particularly infection of mice with the natural whipworm *Trichuris muris*, which is genotypically and phenotypically similar to *T. trichiura* (Klementowicz et al., 2012; Else et al., 2020). Despite decades of research using *T. muris*, fundamental aspects of *Trichuris* development remain poorly understood. Foundational descriptions of *T. muris* morphology are outdated and largely schematic, with limited quantitative detail on growth dynamics across the full length of the parasite life cycle, especially within whole-body context (Fahmy, 1954; Wakelin, 1969; Panesar, 1989; Blasco-Costa and Poulin, 2017; Panti-May et al., 2023; Airs and Duque-Correa, 2025).

Progress in understanding whipworm biology has been further constrained by the lack of robust *in vitro* culture systems that support parasite development. Consequently, studies have relied almost exclusively on *in vivo* mouse infections, which limit experimental control and mechanistic dissection of host–parasite interactions at the epithelial interface. Whipworm development occurs in the context of complex and sustained interactions with host IECs. This prolonged and intimate intracellular interaction is unique among gastrointestinal nematodes and suggests that the interactions between the parasite and host IECs play a central role in regulating whipworm development and persistence. Indeed, larvae hatched *in vitro* in the absence of host cells fail to grow or moult and become developmentally arrested, highlighting the essential role of host-derived cues.

Here, we have pioneered the development of caecal organoids (termed “caecaloids”) as a system to support the *in vitro* growth and development of *T. muris* larvae. Caecaloids closely recapitulate the cellular composition, spatial architecture, and biochemical environment of the caecal epithelium (Duque-Correa et al., 2020). Our previous work using caecaloids showed successful invasion and early stages of infection by T. *muris* L1 larvae over the first 72 hours of infection (Duque-Correa et al., 2020; Duque-Correa et al., 2022). In this study, we demonstrate that caecaloids support the growth and morphological development of *T. muris* larvae for extended periods, where larvae successfully progressed through L2 and L3 stages, reaching substantial size and a high level of morphological complexity and recapitulating their development *in vivo*. Our work thereby provides validation for caecaloids as a physiologically relevant model for studying whipworm infection and development and establish a foundation for future mechanistic studies of host–whipworm interactions.

## Materials and Methods

### Mice

NOD.Cg-*Prkdc^scid^ Il2rg^tm1Wjl^*/SzJ (NSG) mice were maintained under pathogen-free conditions. All mice were housed under a 12-h light/dark cycle at a temperature of 19–24°C and 40–65% humidity. Mice were fed a regular autoclaved chow diet (LabDiet) and had *ad libitum* access to food and water. All efforts were made to minimise suffering by considerate housing and husbandry. Animal welfare was assessed routinely for all mice involved. Mice were naïve prior to the studies here described. Experiments were performed under the regulation of the UK Animals Scientific Procedures Act 1986 under the Project licence PP3418746 and were approved by the University of Cambridge Animal Welfare and Ethical Review Body.

### Trichuris muris life cycle

Infection and maintenance of *T. muris* was conducted as described (Wakelin 1967). Briefly, NSG mice (7-29 weeks old) were orally infected with a high (n ≈ 400 eggs) dose of embryonated eggs from *T. muris* Edinburgh (E)-isolate. Mice were monitored daily for general condition and weight loss. For life cycle maintenance, thirty-five days later, mice were culled by cervical dislocation and the caecum and proximal colon were removed. The caecum was split and washed in RPMI-1640 plus 500 U/mL penicillin and 500 μg/mL streptomycin (all from Gibco Thermo Fisher Scientific). Worms were removed using fine forceps and cultured for 4 h or overnight in RPMI-1640 plus 500 U/mL penicillin and 500 μg/mL streptomycin at 37°C, 5% CO_2_. Media where the worms were cultured, containing eggs and excretory/secretory products was centrifuged at 720 *g*, for 10 min, at 21–22°C (room temperature, RT), with no brake, to pellet the eggs. The eggs were allowed to embryonate for eight weeks in distilled water in the dark at RT (White et al. 2018; Duque-Correa et al. 2019). Upon completion of embryonation, eggs were long-term stored at 4°C, and infectivity was determined based on worm burdens in NSG mice infected with a high dose of eggs and culled at day 35 post infection (p.i.).

### Recovery of *T. muris* larval and adult stages from infected mice

*Trichuris muris*-infected NSG mice were culled by cervical dislocation after 7, 14, 20, 25 and 35 days p.i. to recover L1, L2, L3 and L4 larvae and adults, respectively. Caecum and proximal colon were collected, placed in Dulbecco’s PBS 1X without calcium and magnesium (PBS) containing 500 U/mL penicillin and 500 μg/mL streptomycin (all from Gibco Thermo Fisher Scientific), cut longitudinally and washed to remove faecal contents. For L1-L3 larvae recovery, the tissues were cut into small sections and added to 0.9% NaCl (Merck) in PBS, and incubated in a water bath at 37°C for 2 h with vigorous shaking every 30 min to release L1-L3 larvae from the epithelium. Larvae were collected from the NaCl solution through inspection under a light microscope and placed into PBS plus 100 U/mL penicillin and 100 μg/mL streptomycin. L4 larvae and adults were removed using fine forceps and placed into PBS plus 100 U/mL penicillin and 100 μg/mL streptomycin.

### *In vitro* hatching of *T. muris* eggs with *Escherichia coli* and recovery of freshly hatched L1 larvae

*Escherichia coli* K-12 was grown in Luria Bertani broth (Becton Dickinson Difco) overnight at 37°C and shaking at 200 rpm. *T. muris* eggs were added to bacterial cultures and incubated for 2 h at 37°C, 5% CO_2_. Larvae were washed with PBS three times to remove *E. coli* by centrifugation at 720 *g* for 10 min at RT brake 3. Bacteria were killed by culturing larvae in complete RPMI-1640 containing 10% Foetal Bovine Serum (FBS) (Gibco Thermo Fisher Scientific), 2 mM Glutamax (Gibco Thermo Fisher Scientific), 1X antibiotic/antimycotic (Sigma-Aldrich) and 1 mg/mL ampicillin (Roche) for 2 h at 37°C, 5% CO_2_. Larvae were washed with complete RPMI-1640 three times to remove ampicillin and separated from egg shells and unembryonated eggs using a stepped 50-60% Percoll (Sigma-Aldrich) gradient. Centrifugation at 300 *g* for 15 min at RT with no brake was performed and the 50% interface layer was collected. Recovered larvae were washed with complete RPMI-1640 and resuspended in media containing 100 μg/mL Primocin (InvivoGen).

### 3D caecaloid culture

Mouse 3D caecaloids lines from C57BL/6N adult mice (6-8 weeks old) were derived from caecal epithelial crypts as previously described (Duque-Correa et al. 2020). Briefly, the caecum was cut open longitudinally and luminal contents removed. Tissue was then cut in small segments that were washed with ice-cold PBS and vigorous shaking to remove mucus, and treated with Gentle Cell Dissociation Reagent (STEMCELL Tech) for 15 min at RT with continuous rocking. Released crypts were collected by centrifugation, washed with ice-cold PBS, resuspended in 200 μL of cold Matrigel (Corning), plated in 6-well tissue culture plates and overlaid with a Wnt-rich medium containing base growth medium (Advanced DMEM/F12 with 2 mM Glutamax, 10 mM HEPES, 1X penicillin/streptomycin, 1X B27 supplement, 1X N2 supplement (all from Gibco Thermo Fisher Scientific)), 50% Wnt3a-conditioned medium (Wnt3a cell line, kindly provided by the Clevers laboratory, Utrecht University, Netherlands), 10% R-spondin1 conditioned medium (293T-HA-Rspo1-Fc cell line, Trevigen), 1 mM N-acetylcysteine (Sigma-Aldrich), 50 ng/mL rmEGF (Gibco Thermo Fisher Scientific, Qkine), 100 ng/mL rmNoggin (Peprotech, Qkine), 100 ng/mL rhFGF-10 (Peprotech, Qkine) and 10 μM Rho kinase (ROCK) inhibitor (Y-27632) dihydrochloride monohydrate (Abcam). Caecaloids were cultured at 37°C, 5% CO_2_. The medium was changed every two days and after one week, Wnt3a-conditioned medium was reduced to 30% and penicillin/streptomycin was removed (expansion medium). Expanding caecaloids were passaged after recovering from Matrigel using Cell Recovery Solution (Corning), by physical dissociation through vigorous pipetting with a p200 pipette every six to seven days.

### Caecaloid culture in 2D conformation using transwells

Two dimensional (2D) caecaloids cultures in transwells were prepared as previously described (Duque-Correa et al. 2022). 3D caecaloids grown in expansion medium for 4-5 days after passaging were dissociated into single cells by TrypLE Express (Gibco Thermo Fisher Scientific) digestion. Two hundred thousand cells in 200 µL base growth medium were seeded onto 12 mm transwells with polycarbonate porous membranes of 0.4 µm (Corning) pre-coated with 50 mg/mL rat tail collagen I (Gibco Thermo Fisher Scientific). Cells were cultured with expansion medium in the basolateral compartment for two days. Then, basolateral medium was replaced with medium containing 10% Wnt3a-conditioned medium for an additional 48 h. To induce differentiation of cultures, medium in the apical compartment was replaced with 50 μL base growth medium and medium in the basolateral compartment with medium containing 2.5% Wnt3A-conditioned medium that was changed every two days. Cultures were completely differentiated when cells pumped the media from the apical compartment and cultures looked dry.

### *Trichuris muris* L1 larvae infection of caecaloids grown in transwells

Differentiated caecaloid cultures in transwells were infected with 300 L1 *T. muris* larvae obtained by *in vitro* hatching of eggs in presence of *E. coli*. Larvae in a volume of 100 μL of base growth medium were added to the apical compartment of the transwells. Infections were maintained for up to 20 days at 37°C, 5% CO_2_. Throughout culture, medium in the basolateral compartment was replaced every two days with fresh medium containing 2.5% Wnt3A-conditioned medium.

### Immunofluorescence (IF) staining of parasites

Isolated life stages of *T. muris* were fixed in 4% formaldehyde, methanol-free (Thermo Fisher) in PBS for 20 min to 1 h on rotation at RT, washed three times in PBS and stored at 4°C. For staining, samples were permeabilised in 2% Triton-X100 (Sigma-Aldrich) and 5% FBS in PBS for 1 h on rotation at RT, followed by incubation with 4’,6’-diamidino-2-phenylindole (DAPI, 1:1000, AppliChem, A1001.0010) and phalloidin Alexa Fluor 488 (1:1000, Invitrogen, A12379) or 647 (1:1000, Thermo Fisher Scientific, A22287) in PBS with 0.25% Triton-X100 on rotation at 4°C overnight (∼16 h). Stains were then washed in PBS with 0.05% Tween (Sigma-Aldrich) three times, and individual worms were transferred by wide bore pipette tip or directly to slides, micro-manipulated to separate individuals using fine probes or a p20 pipette, and mounted in Fluoromount G (Invitrogen Thermo Fisher Scientific).

### IF staining of caecaloids

Caecaloid cultures in transwells were fixed with 4% formaldehyde, methanol-free in PBS for 20 min at 4°C, washed three times with PBS and permeabilised with 2% Triton X-100 5% FBS in PBS for 1 h at RT. Caecaloids were then incubated with primary antibodies α-villin (1:100, Abcam, ab130751), α-Ki-67 (1:250, Abcam, ab16667), α-chromogranin A (1:50, Abcam, ab15160), α-Dcamlk-1 (1:200, Abcam, ab31704), and the lectins *Ulex europaeus* agglutinin - Atto488 conjugated (UEA, 1:100, Sigma-Aldrich, 19337) and *Sambucus nigra -* Fluorescein conjugated (SNA, 1:50, Vector Laboratories, FL-1301) diluted in 0.25% Triton X-100 5% FBS in PBS overnight at 4°C. After three washes with PBS, caecaloids were stained with secondary antibody Donkey anti-rabbit IgG Alexa Fluor 555 (1:400, Molecular Probes, A31572), phalloidin Alexa Fluor 647 (1:1000) and DAPI (1:1000) at RT for 1 h. Transwell membranes were washed three times with PBS and mounted on slides using ProLong Gold anti-fade reagent (Life Technologies Thermo Fisher Scientific).

### Imaging and morphobiometric data measurements

Imaging of fixed and stained *in vivo* specimens was conducted on: 1) a Zeiss Axiovert 5 for brightfield length measurements; 2) a Leica DMI8 Thunder Imager to acquire representative phase contrast and immunofluorescence images of all life stages and additional length and tissue measurements of L1 larvae at day 7p.i. worms; and 3) a Leica SP8 confocal microscope, for tissue relative proportion and cell segmentation analyses of worms at other life stages. Imaging of fixed and stained caecaloid culture was conducted on a Leica SP8 confocal microscope, for additional length and tissue measurements of larvae at day 7 and 20 p.i. Presence of structures in *in vitro* larvae was assessed during live acquisition on a Leica SP8 confocal microscope.

Brightfield *in vivo* images with multiple frames were stitched using the Pairwise Stitching tool in ImageJ, and worm lengths were measured using the Segmented Line tool in ImageJ. Fluorescent and confocal microscopy images were processed using the Leica Application Suite X software. Whole-body and substructure length were measured from x-y projected z-stack or individual confocal planes, depending on the structure of interest, using built-in tools in Zeiss Arivis Pro software.

Substructures measured (Table 1) were decided based on parameters reported by (Panti-May et al. 2023). Morphological identification was conducted based on features described in *T. suis* by (Beer 1973) and *T. trichiura* by (Rivero et al. 2020).

**Table 1.** Main morphological features of *Trichuris muris* and corresponding analysis methods.

| Feature | Start point | Start channel | Stop point | Stop channel | Measurement axis | Used *in vitro*? |
| --- | --- | --- | --- | --- | --- | --- |
| **Whole body** | Tip of head | Brightfield | End of body | Brightfield | Midline of body | Yes but with DAPI and Phalloidin as start and stop channels |
| **Tip of head to nerve ring** | Tip of head | Brightfield / Phalloidin | Middle of nerve ring | DAPI | Midline of body | Yes, DAPI and phalloidin start and stop |
| **Oesophagous to Stichosome entrance** | Tip of head | Brightfield / Phalloidin | Stichosome start / End of oesophagus (day 0 and day 7 worms) | DAPI / Phalloidin / Brightfield | Midline of tissue | Yes, DAPI and phalloidin start and stop |
| **Stichosome** | Start of stichosome | DAPI / Phalloidin / Brightfield | End of stichosome / start of intestine | DAPI / Phalloidin / Brightfield | Midline of tissue | Yes, DAPI and phalloidin start and stop |
| **Intestine** | End of stichosome / start of intestine | DAPI / Phalloidin / Brightfield | End of intestine / Start of rectum or cloaca | DAPI / Brightfield | Midline of tissue | Yes but with DAPI and Phalloidin as start and stop channels |
| **Ovary** | Start of ovary | Brightfield / DAPI | End of ovary | Brightfield / DAPI | Midline of tissue | No |
| **Testis** | Start of testis | Brightfield / DAPI | End of testis | Brightfield / DAPI | Midline of tissue | No |
| **Rectum/Cloaca** | Start of rectum or cloaca | Brightfield / DAPI | End of body | Brightfield | Midline of tissue | Yes but with DAPI and Phalloidin as start and stop channels |

### Statistical analyses

Statistical comparisons for biometrical data of worms from two different age groups were performed using Mann–Whitney U two-tailed tests from the Prism 10 software (GraphPad).

## Results

### Trajectory of *T. muris* growth is independent of host mouse strain

*Trichuris muris* growth and moulting through its life cycle was described by Panesar three decades ago (Panesar, 1989). Since then, these observations have not been revisited or cross-compared across experimental systems. Moreover, developmental dynamics of *T. muris* may vary depending on the host mouse strain, which can influence parasite growth and maturation. To establish an up-to-date reference dataset for validating parasite development in caecaloids, we compared the growth, based on total body length, of isolated fixed individual worms of the Edinburgh strain raised in NSG mice to previous descriptions of the same strain raised in DBA/2 (Panesar, 1989) and outbred albino Schofield and Evans mice (Wakelin, 1969). We measured the whole-body growth of *T. muris* from hatching (L1, day 0 post-infection, p.i.) to adulthood (male and females, day 35 p.i.) in NSG mice at days 7 (L1), 13 (L2), 14 (L2), 20 (L3), 21 (L3), and 25 (L4) p.i. Consistent with observations by both Panesar (1989) and Wakelin (1969), we observed a slow growth rate during the first phase of development (days 0 – 7 p.i.), followed by a marked increase in growth (whole-body length) from day 7 to 14 p.i., which coincides with the first moulting from L1 to L2 taking place between days 9 and 11 p.i.. Beyond 14 days p.i., *T. muris* infecting NSG mice exhibited approximately exponential growth, with the most substantial increase observed between day 25 larvae (L4) and adults (Figs 1 and 2A). This trend was also reported by Panesar (1989) (Fig 1). At 20 to 25 days p.i., Panesar recorded L3 and L4 larvae that are longer than those from our study. This reversed by day 35 p.i., at which point both male and female adult worms in our study were slightly longer than those recorded by Panesar (Fig 1). In our study, at day 35 p.i., adult female worms were significantly longer than males (Figs 1 and 2A); this was not the case between adult female and male worms at days 35 and 40 p.i. in Panesar’s study (1989). Taken together, while we observed subtle variations in *T. muris* growth in different mouse strains, our results indicate that the overall pattern of *T. muris* whole-body growth from hatching to adulthood is largely conserved and independent of host mouse genetic background.

**Figure 1.**
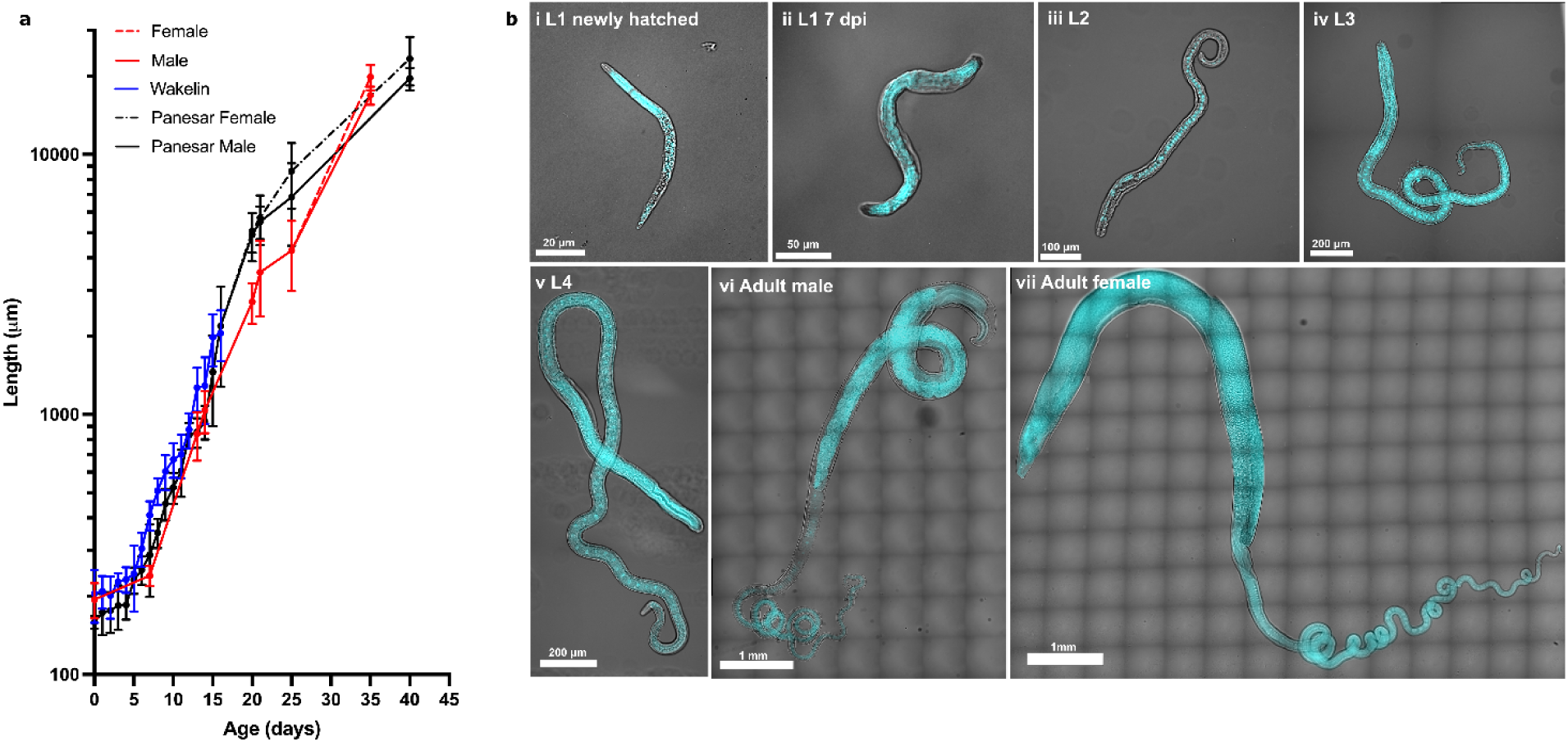
Trajectory of *Trichuris muris* growth in different host mouse strains. **(a)** Mean whole-body length (logarithmic scale) plotted against age (arithmetic scale) of *T. muris* during development in DBA/2 (black line) (Panesar, 1989), outbred albino Schofield and Evans (blue line) (Wakelin, 1969) and NSG (red line) mice (this study). Means with standard deviation are shown. Mean length at day 0 in Panesar and this study is based on measurements of newly hatched larvae *in vitro*, while Wakelin collected larvae 2 h post infection (p.i). Panesar measured the total body length of *T. muris* every day until day 40 p.i. (n ∼ 20). Wakelin measured the total body length of *T. muris* every day until day 16 p.i. (n > 25). In this study, total length of *T. muris* recovered at days 0 (n = 84), 7 (n = 35), 13 (n = 29), 14 (n = 51), 20 (n = 48), 21 (n = 49), 25 (n = 39), 35 (females n = 13 and males n = 22) p.i. was measured in brightfield images using ImageJ. **(b)** Representative images showing (i) *T. muris* first-stage (L1) larvae hatched *in vitro* and gross *in vivo* growth and development of *T. muris* recovered from *T. muris*-infected NSG mice at days (ii) 7, (ii) 14, (iii) 20, (iv) 25, and (vi-vii) 35 post infection. A single focal plane from each z-stack is shown to facilitate visualisation of internal structures. Phase images are overlaid with DAPI (cyan, labelling nuclei).

**Figure 2.**
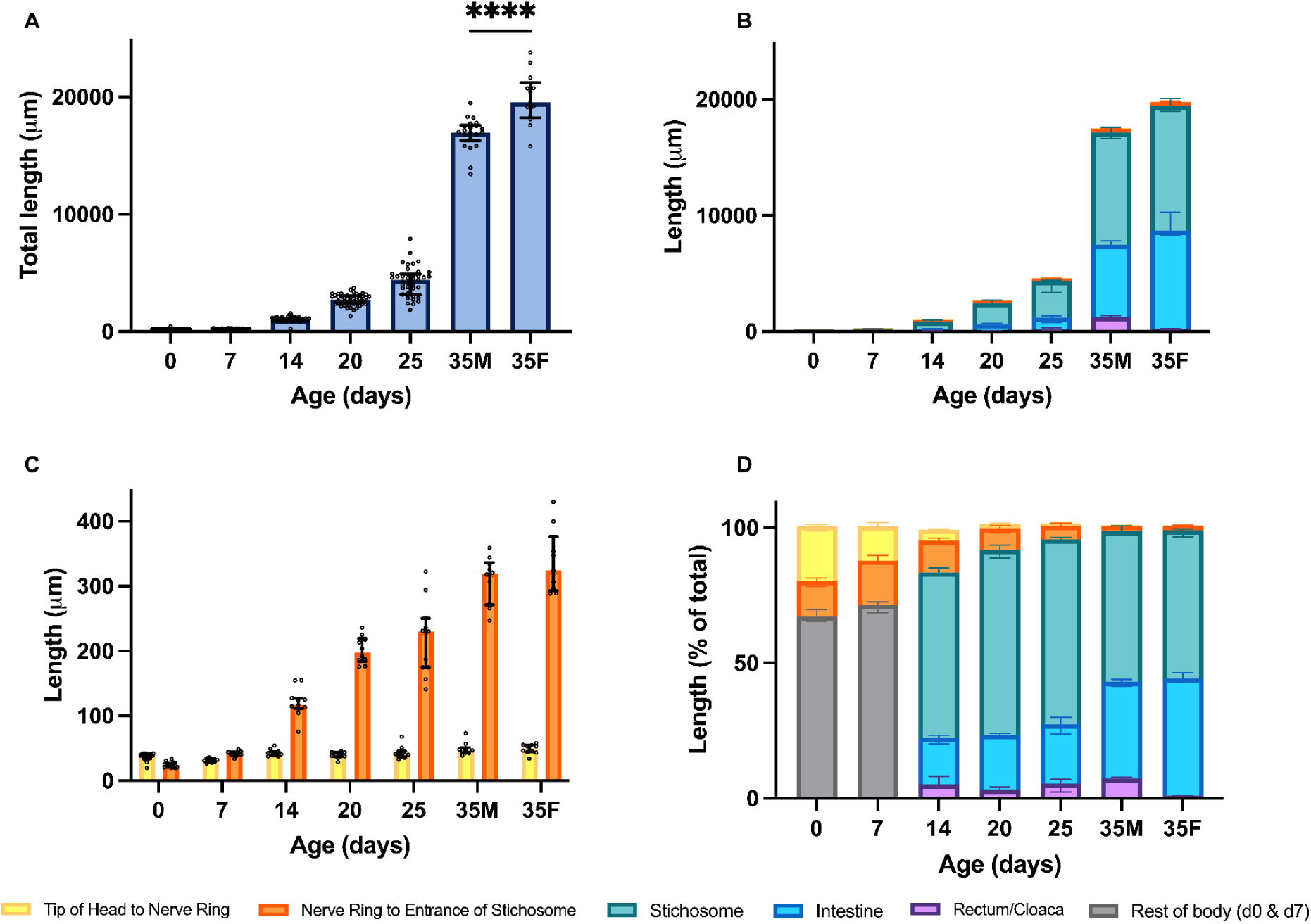
Biometrical data of *Trichuris muris* across *in vivo* development. *Trichuris muris* first-stage (L1) larvae hatched in vitro and second to fourth (L2-L4) larvae and adult worms collected from NSG infected mice were stained with DAPI and phalloidin to visualised nuclei and F-actin, respectively. Z-stack confocal images were acquired and total and substructure length were measured using Zeiss arivis Pro software. (A) Total body length of *T. muris* recovered at days 0 (n = 84), 7 (n = 35), 14 (n = 51), 20 (n = 48), 25 (n = 39), and 35 (females n = 13 and males n = 22) p.i.. Median with individual data points and interquartile range are shown. ****p < 0.001 by Mann-Whitney test. (B) Length of tip of head to nerve ring (NR), length from NR to entrance of stichosome (or to end of oesophagus for day 0 and day 7), stichosome (ST), intestine (IN), rectum/cloaca (REC/CL) were measured on *T. muris* at days 0 (n_NR_ = 17, n_NR to ST_ = 14, n_Rest of body_ = 17), 7 (n_NR_ = 10, n_NR to ST_ = 9, n_Rest of body_ = 10), 14 (n_NR_ = 11, n_NR to ST_ = 11, n_ST_ = 11, n_IN_ = 11, n_REC_ = 11), 20 (n_NR_ = 10, n_NR to ST_ = 10, n_ST_ = 10, n_IN_ = 10, n_REC_ = 10), 25 (n_NR_ = 10, n_NR to ST_ = 11, n_ST_ = 12, n_IN_ = 12, n_REC_ = 12), and 35 (females (n_NR_ = 10, n_NR to ST_ = 9, n_ST_ = 10, n_IN_ = 10, n_REC_ = 10) and males (n_NR_ = 10, n_NR to ST_ = 10, n_ST_ = 10, n_IN_ = 10, n_CL_ = 10) p.i.. Median with interquartile range are shown. (C) Length from tip of head to nerve ring (NR) and from NR to entrance of stichosome (ST) of *T. muris* at days 0 (n_NR_ = 31, n_NR to ST_ = 14), 7 (n_NR_ = 10, n_NR to ST_ = 9), 14 (n_NR_ = 11, n_NR to ST_ = 11), 20 (n_NR_ = 10, n_NR to ST_ = 10), 25 (n_NR_ = 10, n_NR to ST_ = 11), and 35 (females (n_NR_ = 10, n_NR to ST_ = 9) and males (n_NR_ = 10, n_NR to ST_ = 10)) p.i.. Median with individual data points and interquartile range are shown. (D) Tissue lengths expressed as a percentage of total body length from *T. muris* recovered at days 0 (n_NR_ = 14, n_NR to ST_ = 14, n_Rest of body_ = 14), 7 (n_NR_ = 10, n_NR to ST_ = 9, n_Rest of body_ = 9), 14 (n_NR_ = 11, n_NR to ST_ = 11, n_ST_ = 11, n_IN_ = 11, n_REC_ = 11), 20 (n_NR_ = 10, n_NR to ST_ = 10, n_ST_ = 10, n_IN_ = 10, n_REC_ = 10), 25 (n_NR_ = 10, n_NR to ST_ = 11, n_ST_ = 11, n_IN_ = 11, n_REC_ = 11),, and 35 (females (n_NR_ = 10, n_NR to ST_ = 9, n_ST_ = 10, n_IN_ = 10, n_REC_ = 10) and males (n_NR_ = 10, n_NR to ST_ = 10, n_ST_ = 10, n_IN_ = 10, n_CL_ = 10) p.i.. Median with interquartile range are shown.

### *in vivo T. muris* development at the tissue level

Throughout their life cycle, whipworms develop from relatively simple L1 larvae, where only a stylet, nerve ring and oesophagus are visible, into complex animals, with adult worms exhibiting sophisticated organs such as the stichosome, intestine and reproductive female and male organs (Fahmy, 1954; Beer, 1973). While camera lucida drawings illustrating *T. suis* (pig whipworms) morphology through its life cycle are available (Beer, 1973), the anatomical descriptions for *T. muris* remain fragmented. Moreover, much of the existing morphological information is not accompanied with microscopic images, or is isolated from the parasite whole-body context, making it difficult to interpret developmental progression. To address these limitations and to generate an updated anatomical reference framework for validating parasite development in caecaloids, we sought to produce a detailed anatomical and morphometric characterisation of *T. muris* development. For this, we acquired high-resolution fluorescent and confocal images of whole worms and quantitatively measured the lengths of major anatomical substructures across developmental stages at day 0 (L1 larvae newly hatched *in vitro*), and days 7 (L1), 14 (L2), 20 (L3), 25 (L4) and 35 (adult) post infection (p.i.) (Figs 1, 2 and 3; Supplementary Fig 1).

**Figure 3.**
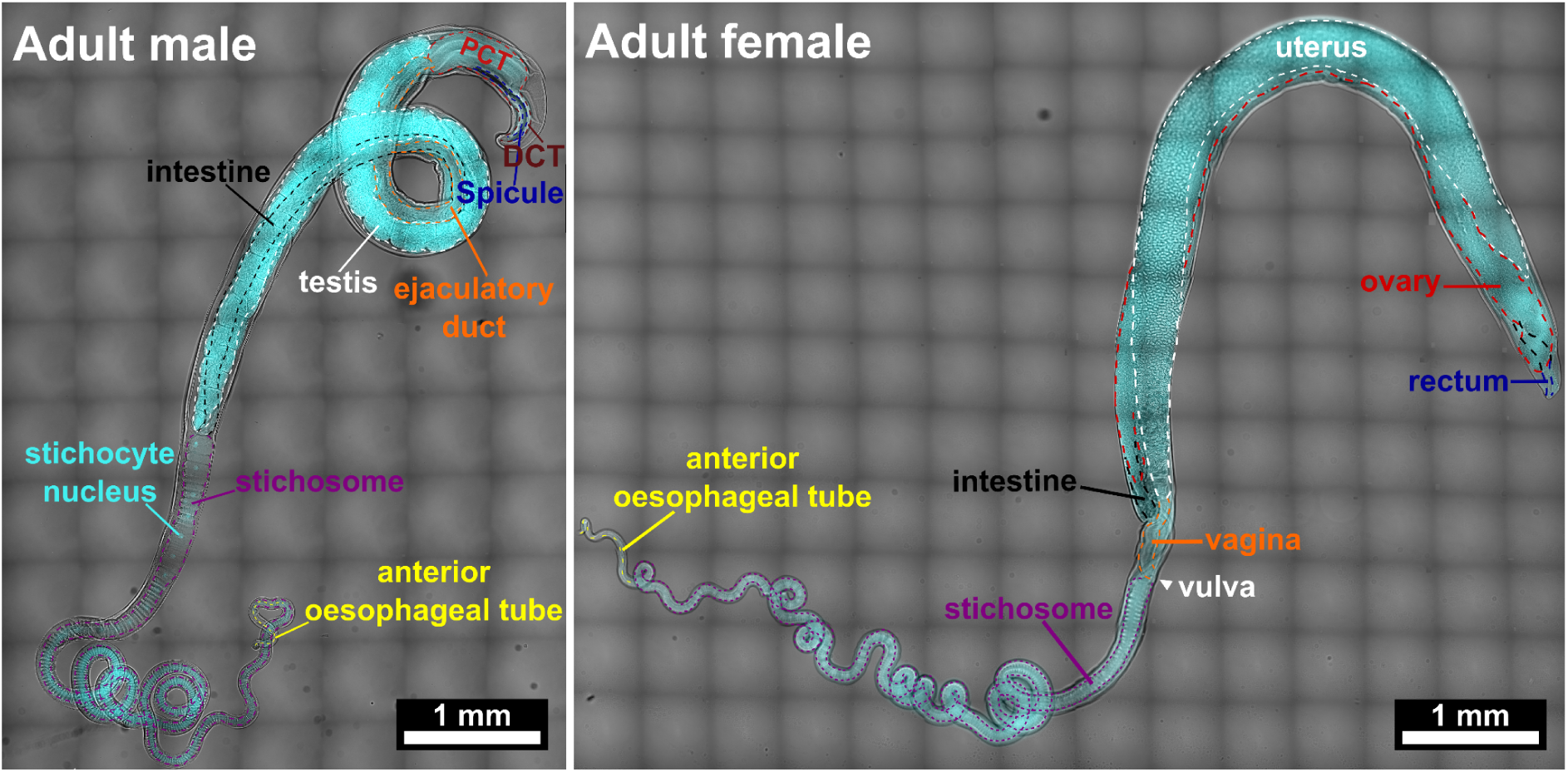
Morphological complexity of adult *Trichuris muris* recovered from *in vivo* infections. Representative images showing morphological complexity and sexual dimorphism of adult male and female *Trichuris muris* recovered from NSG mice infected with *T. muris* eggs. A single focal plane from each z-stack is shown to facilitate visualisation of internal structures. Phase images are overlaid with DAPI (cyan, labelling nuclei). Dashed lines indicate internal organ boundaries. PCT, proximal cloacal tube; DCT, distal cloacal tube.

Consistent with previous reports, in L1 larvae recovered at days 0 and 7 p.i., the nerve ring and oesophagus were the only noticeable internal structures (Figs 1 b). At day 14 p.i., L2 larvae showed a major increase in anatomical complexity, where a thicker and more convoluted oesophagus, a prominent stichosome, a clear intestine, and a subterminal rectum that leads to an anus were clearly visible (Fig 1b). At days 20 and 25 p.i., L3 and L4 larvae continue to grow substantially in size (Fig 1 a & 2 a) and start to exhibit clearer sexual dimorphism. The worms reached adulthood at day 35 p.i., with complete morphological maturity (Fig 3).

Throughout *T. muris* life cycle, the nerve ring was distinguished as a distinct band lacking nuclei with no DAPI fluorescence staining (Fig 1b). The oesophagus was visible as a thin continuous tube extending from the mouth of the worm. In newly hatched larvae (L1, day 0), the oesophagus drained into a hollow body cavity, where nuclei density was lower around the oesophageal tube and higher over the hollow body cavity (Fig 1 bi & ii). As the worms develop, the oesophagus appeared much more convoluted and became intracellular as it enters and runs within the stichosome. Notably, while the anterior region of *T. muris* body increased significantly in length as the worms develop, the length from the tip of the head to the nerve ring increased only minimally, and most growth of the anterior body occurred posterior to the nerve ring (Fig 2 c). The anterior body region forms the characteristic slender part that remains embedded within the syncytial tunnel of host IECs at the L3 to adult stages, and is made up mainly of the stichosome surrounded by the outer cuticle and bacillary band. From the point of its emergence (L2) through to adulthood, the stichosome stood out as one of the most prominent structures, taking up more than half of the worm body (Fig 2 b & d). Notably, although most L2 larvae possessed stichosomes that compose of a single row of large stichocytes, some worms exhibited a double-row arrangement of stichocytes toward the posterior region of the stichosome, suggesting that ongoing development of this structure may occur during the L2 stage (data not shown).

Coinciding with the end of the stichosome is the oesophagus-intestinal junction, marking also the transition from the anterior to posterior region of *T. muris* body. The posterior body contains the intestine and reproductive organs. While throughout larval stages (L2 to L4), there is no significant difference in the thickness of the anterior and posterior body regions, the posterior body region of adult worms are visibly thicker than their anterior body region (Fig 1b), likely reflecting the need to accommodate the highly developed reproductive organs. While the intestinal lumen could be clearly observed at the L2 stage (Fig 1b), visualisation became more challenging at later larval and adult stages, as the intestine was partially obscured by the developing reproductive organs that run alongside it. At these stages, the intestine was visible as a thin muscular tube (Fig 3).

While it was difficult to visualise the whole length of the developing reproductive structures to obtain quantitative biometrical measurements, we observed the same developmental progression of the female reproductive system as described by Beer (1973) in *T. suis*. Specifically, at day 14 p.i. (L2), the female reproductive system was a simple strand that is relatively straight with a thicker proximal region starting close to the oesophageal-intestinal junction (data not shown). At day 20 p.i. (L3), the slender oviduct connecting the uterus and ovary began to extend and formed into a loop. The vulva opening appeared as a thickened cuticular swelling close to the oesophageal-intestinal junction (Fig 3) from day 20 p.i. (L3) through to adulthood. In both male and female adult worms, the reproductive systems are highly developed and sophisticated structures that take up a large portion of the worm body (Supplementary Fig 1). The adult female reproductive system is a highly convoluted and complex structure. The uterus spans the whole length of the posterior body and appeared darker and expanded from being packed full of eggs (Fig 3). During egg laying, adult female worms displayed rhythmic contractions that appeared to propel the eggs from the uterus through the short narrow vagina and out of the vulval opening. At the distal end of the uterus, a thin narrow oviduct forms a subterminal loop that connects the uterus to the ovary. The ovary appeared as thick darky granulated muscular tube that also spans the whole length of the posterior body (Fig 3). On the other hand, the adult male reproductive system was relatively less sophisticated, with the testis resembling the ovary in appearance (thick, darkly granulated muscular tube) and also spanning the whole length of the worm’s posterior body (Fig 3). The seminal vesicle and vas deferens that are described by Beer (1973) to connect the testis with the ejaculatory duct in *T. suis* were difficult to observe. We did, however, observe the ejaculatory duct as a tubular structure with a ridged surface that runs in parallel with the posterior part of the intestine, both of which appeared to join into the proximal cloacal tube (PCT) (Fig 3). The cloaca in adult male *T. muris* is significantly longer than the rectum in adult female worms, consistent with previous observations in *T. suis* (Beer, 1973) (Fig 2 b & d; Supplementary Fig 1). While the female adult rectum is a subterminal short simple tubular structure that opens to the anus, the adult male cloaca is made up of two parts – the PCT and the distal cloacal tube (DCT). The PCT is a wide tubular structure with a large lumen that is continuous with the DCT. The DCT is narrower, containing the spicule and leading to the cloacal opening (Fig 3).

Together, these novel microscopy and biometrical data provide a detailed and contextualised description of the growth and morphological development of *T. muris,* providing a valuable resource to be used as reference for the evaluation of *T. muris* larvae obtained from *in vitro* infection of caecaloids as well as for future anatomical studies.

### Caecaloids support *T. muris* growth and development *in vitro*

The molecular and cellular cues triggering and sustaining whipworm development remain unknown. Due to the intracellular lifestyle of *Trichuris* spp within the intestinal epithelial cells, we hypothesise that physicochemical interactions of the parasite with the caecal epithelium provide critical signals for whipworm growth and moulting. Our laboratory has successfully reproduced *T. muris* invasion and the formation of syncytial tunnels within the intestinal epithelia using caecaloids. This *in vitro* model has enabled us to characterise the early events (first three days) of infection by L1 larvae (Duque-Correa et al., 2022). To investigate if the caecal epithelium is sufficient to sustain whipworm development, we extended our caecaloid-*T. muris* co-cultures and evaluated parasite growth and morphogenesis after 1, 3, 7, 11, 13, and 20 days p.i. (Figs 4 - 8).

**Figure 4.**
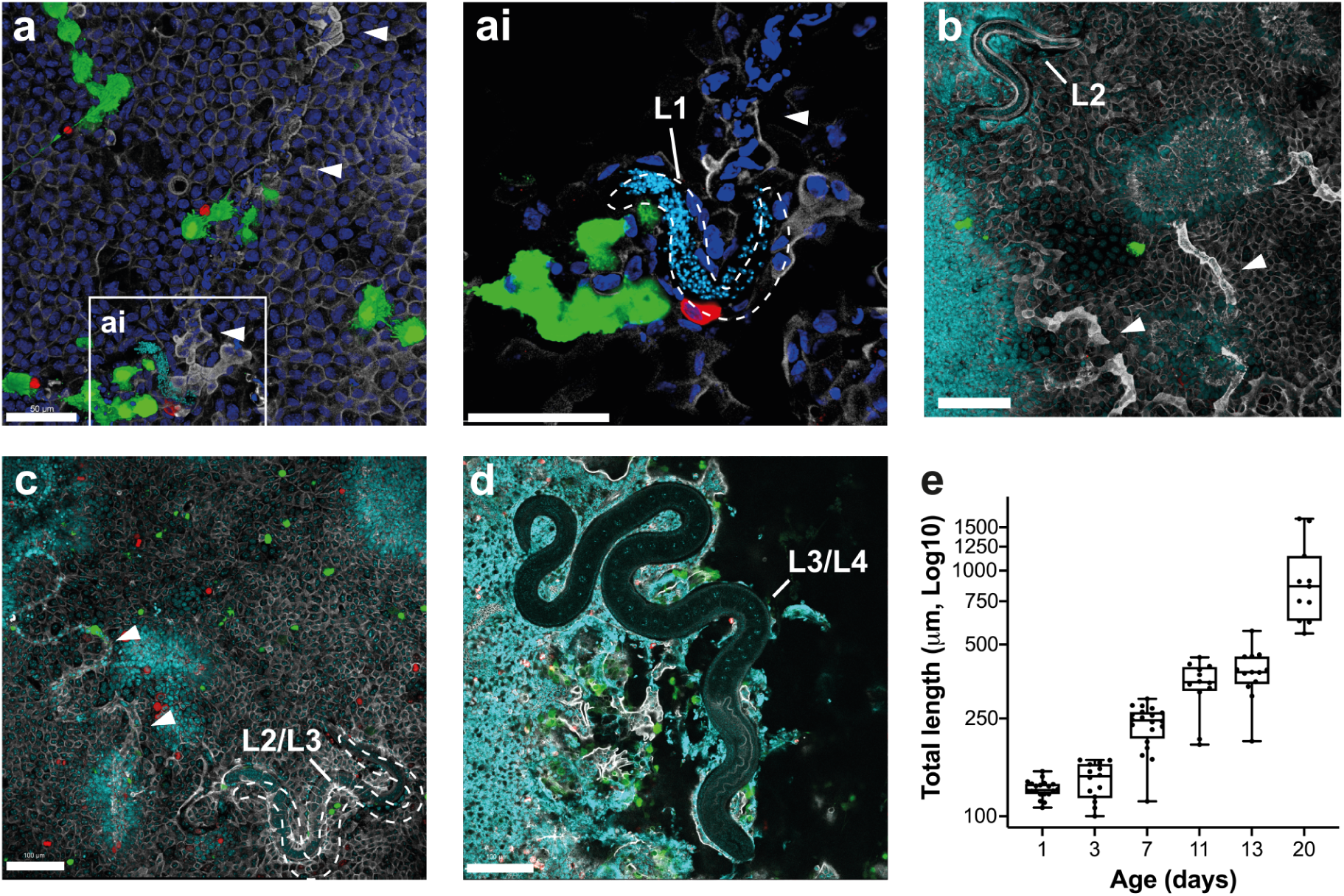
Caecaloids support *Trichuris muris* growth *in vitro.* Representative confocal immunofluorescence (IF) images of *Trichuris muris**-***infected caecaloids showing larval growth from (a-ai) day 1, (b) day 13, and (c-d) day 20 post infection. Scale bars for (a & ai) 50 µm and (b-d) 100 µm. Arrow heads show syncytial tunnels. Dashed lines show larval boundaries. Complete z-stack projections showing larvae infecting IECs. In green, the lectins UEA and SNA bind mucins in goblet cells; in red, (a) Dclk-1, marker of tuft cells, (b-d) Ki-67 labels dividing cells; in blue and aqua, DAPI stains nuclei of IECs and larvae; and in white, phalloidin binds to F-actin. (e) Length of larvae over time in infected caecaloids at days 1 (n = 20), 3 (n = 14), 7 (n = 20), 11 (n = 12), 13 (n = 13), and 20 (n = 11) p.i. IF imaging experiments on *T. muris-*infected caecaloids were done in triplicate across at least two independent replicas using three caecaloid lines derived from three C57BL/6 mice.

Excitingly, we observed continuous *T. muris* growth over time. While initially slow, with worms at day three p.i only showing slight increase in length compared to day one p.i., after seven days of infection the worms doubled in size (Fig 4 e). From this time point, worms continued to steadily grow, with parasites reaching a maximum whole body length of over 1600 μm at day 20 p.i (Figs 4 c-e, 5 aiii, 7 & 8). Worms measured were those that remained completely intracellular with some actively burrowing and moving within the intestinal epithelial cells, as evidenced by tortuous syncytial tunnels clearly visible in the transwell cultures (Figs 4 a-c, 7 & 8). However, the longer the culture, the more extracellular worms were found, which while alive and moving, did not further grow.

**Figure 5.**
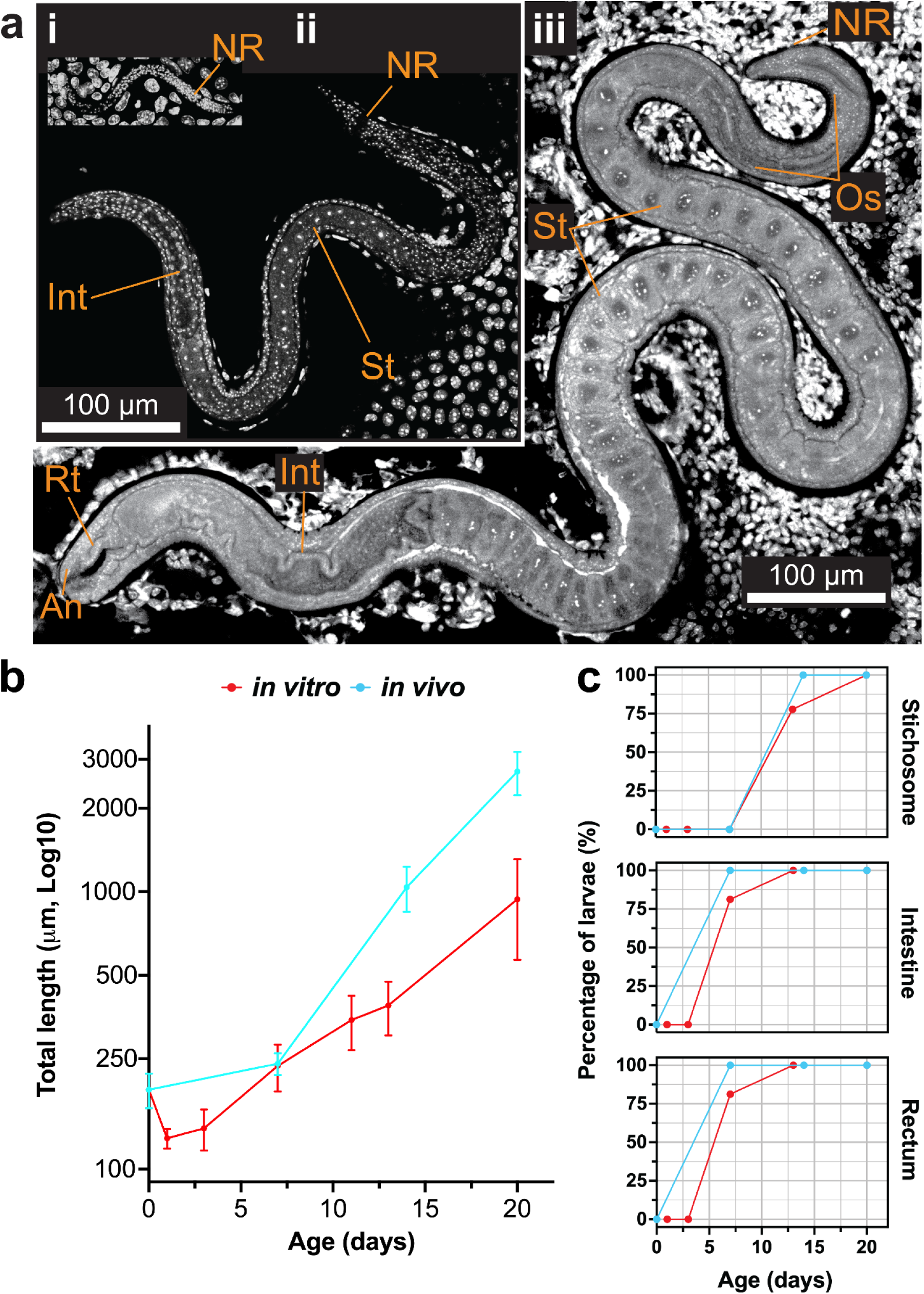
Morphogenesis of *Trichuris muris in vitro* recapitulates *in vivo* development. (a) Confocal microscopy images of representative *T. muris* from infected caecaloids at (ai) L1 stage, (aii) early L2 stage, and (aiii) L3 stage. Structures shown include Os = Oesophagus, NR = Nerve ring, St = Stichocyte in stichosome, Int = Intestine, Rt = Rectum, An = Anus. In white, DAPI stains nuclei of IECs and larvae. (b-c) Comparison of whole body (WB) length (b) and presence of key structures (c) in *T. muris* larvae over time from infections *in vitro* (at days 1 (n_WB_ = 20, n_ST/IN/REC_ = 20), 3 (n_WB_ =14, n_ST/IN/REC_ = 14), 7 (n_WB_ = 20, n_ST/IN/REC_ = 16), 11 (n_WB_ = 12), 13 (n_WB_ = 13, n_ST/IN/REC_ = 11), and 20 (n_WB_ = 11, n_ST/IN/REC_ = 6) p.i.) and *in vivo* (at days 0 (n_WB_ = 84, n_ST/IN/REC_ = 17), 7 (n_WB_ = 35, n_ST/IN/REC_ = 10), 14 (n_WB_ = 51, n_ST/IN/REC_ = 11), and 20 (n_WB_ = 48, n_ST/IN/REC_ = 10) p.i.)

The increase in whole-body length during caecaloid infection was accompanied by an augmentation in the morphological complexity of the parasites (Figs 5 - 8). Larvae in caecaloids infected for one and three days were very simple animals, with the nerve ring and a primitive oesophageal tube being the only evident organs by its lack of nuclei staining. From day 7 p.i., some worms were already showing a developing intestine and primitive stichosome (data not shown). At day 13 p.i., worms showed a more developed stichosome and clear intestine (Fig 5 aii & c, 7a). At day 20 p.i., the larvae have developed into much more complex animals with thicker and more convoluted oesophagi, complete stichosomes with enlarged stichocytes, larger intestines and visible rectums with clear anuses (Figs 4 d, 5 aiii, 7 c, & 8). The larger worms we found at this time point also presented a well-developed bacillary band composed of multiple bacillary cells spread along the cuticle of the parasites (Figure 8c).

Notably, we observed considerable variability in the length and stage of development of worms measured at any time point and indeed in the same caecaloid culture (Figs 4 c-d, & 7 b-c). This highlighted the fact that whipworm development is a continuous and asynchronous process punctuated by periods of growth and moulting (O’Sullivan et al., 2020; Wakelin, 1969). While this made it challenging to define the developmental stage of *in vitro* parasites, this variability presented an unexpected opportunity to capture more stages of whipworm morphogenesis *in vitro*. To determine the developmental stage of the worms *in vitro* as well as to evaluate the robustness of caecaloids in supporting worm growth and development, we obtained a comprehensive biometrical dataset for *in vitro* larvae, examining gross body growth and internal tissue development, and compared with our *in vivo* biometrical dataset (Fig 5 b-c & 6). Excitingly, *in vitro* larvae followed a similar growth trajectory and morphological development pattern compared to larvae retrieved from *in vivo* infections (Fig 5 b-c). Of note, while most *in vitro* larvae exhibited a developing intestine from day 7 p.i., and other specialised structures such as the stichosome and rectum from day 13 p.i. (Fig 5c), it was challenging to obtain exact measurements of these tissue lengths due to technical challenges to visualise these structures within worms that are completely intracellular. At day 20 p.i., the relative proportions of tissue lengths in *in vitro* worms reflect closely those in *in vivo* worms, with the stichosome being one of the longest organs, followed by the intestine (Fig 6c). Within the anterior body region of *in vitro* worms, we also observed that most growth occurs posterior to the nerve ring, whereas the length between the tip of the head and the nerve ring showed only a minor increase (Fig 6a).

**Figure 6.**
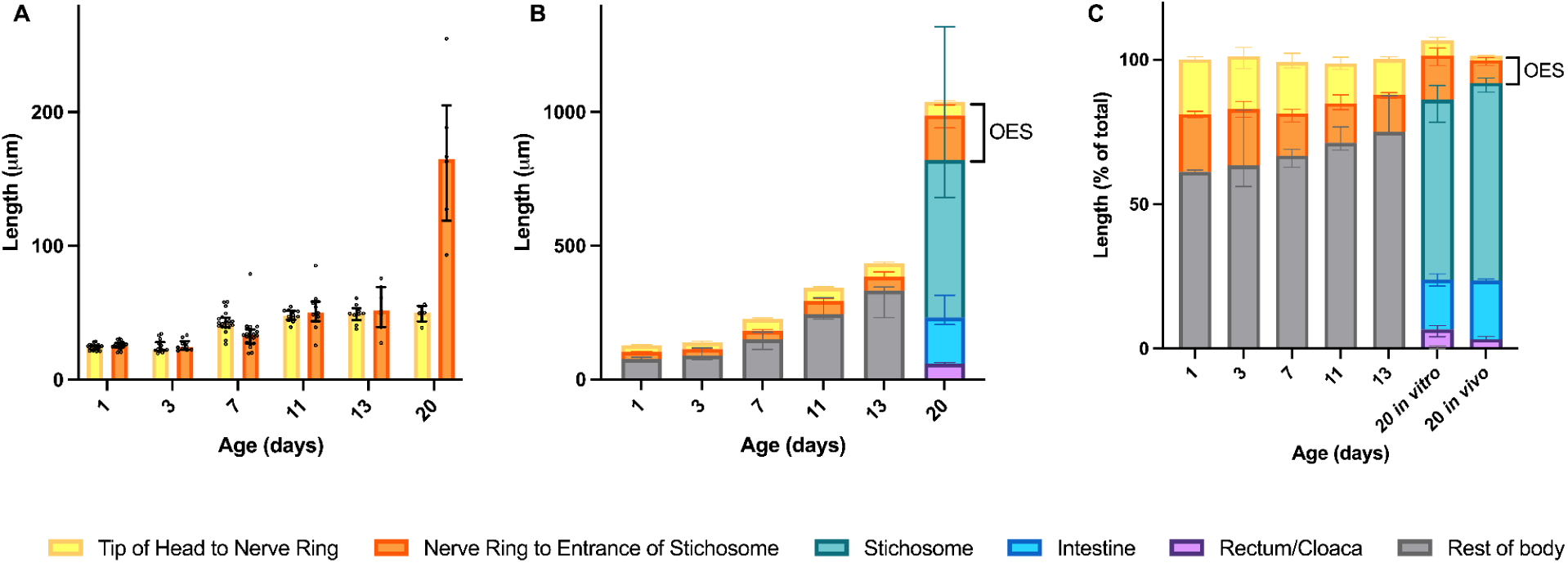
Biometrical data of *Trichuris muris* across *in vitro* development. IF imaging experiments on *T. muris-*infected caecaloids were done in triplicate across at least two independent replicas using three caecaloid lines derived from three C57BL/6 mice. Z-stack confocal images were acquired, and substructure length was measured using Zeiss arivis Pro software. (A) Length from tip of head to nerve ring (NR) and from NR to entrance of stichosome (ST) of *T. muris* recovered from *in vitro* infection of caecaloids at days 1 (n_NR_ = 20, n_NR to ST_ = 20), 3 (n_NR_ = 14, n_NR to ST_ = 10), 7 (n_NR_ = 20, n_NR to ST_ = 20), 11 (n_NR_ = 12, n_NR to ST_ = 11), 13 (n_NR_ = 9, n_NR to ST_ = 7), and 20 (n_NR_ = 6, n_NR to ST_ = 6) p.i.. Median with individual data points and interquartile range are shown. (B) Length of tip of head to nerve ring (NR), length from NR to entrance of stichosome (or to end of oesophagus for days 1 to 13), stichosome (ST), intestine (IN), rectum/cloaca (REC/CL) were measured on *T. muris* recovered from *in vitro* infection of caecaloids at days 1 (n_NR_ = 20, n_NR to ST_ = 20, n_Rest of body_ = 20), 3 (n_NR_ = 14, n_NR to ST_ = 10, n_Rest of body_ = 14), 7 (n_NR_ = 20, n_NR to ST_ = 20, n_Rest of body_ = 15), 11 (n_NR_ = 12, n_NR to ST_ = 11, n_Rest of body_ = 12), 13 (n_NR_ = 9, n_NR to ST_ = 7, n_Rest of body_ = 7), and 20 (n_NR_ = 6, n_NR to ST_ = 6, n_ST_ = 6, n_IN_ = 6, n_REC_ = 5, n_Rest of body_ = 1) p.i.. Median with interquartile range are shown. OES = oesophagus. (C) Tissue lengths expressed as a percentage of total body length from *T. muris* recovered from *in vitro* infection of caecaloids at days 1 (n_NR_ = 20, n_NR to ST_ = 20, n_Rest of body_ = 20), 3 (n_NR_ = 14, n_NR to ST_ = 10, n_Rest of body_ = 14), 7 (n_NR_ = 20, n_NR to ST_ = 20, n_Rest of body_ = 15), 11 (n_NR_ = 12, n_NR to ST_ = 11, n_Rest of body_ = 12), 13 (n_NR_ = 9, n_NR to ST_ = 7, n_Rest of body_ = 7), 20 *in vitro* (n_NR_ = 6, n_NR to ST_ = 6, n_ST_ = 6, n_IN_ = 6, n_REC_ = 5, n_Rest of body_ = 1), and 20 *in vivo* (n_NR_ = 10, n_NR to ST_ = 10, n_ST_ = 10, n_IN_ = 10, n_REC_ = 10) p.i.. Median with interquartile range are shown. OES = oesophagus.

**Figure 7.**
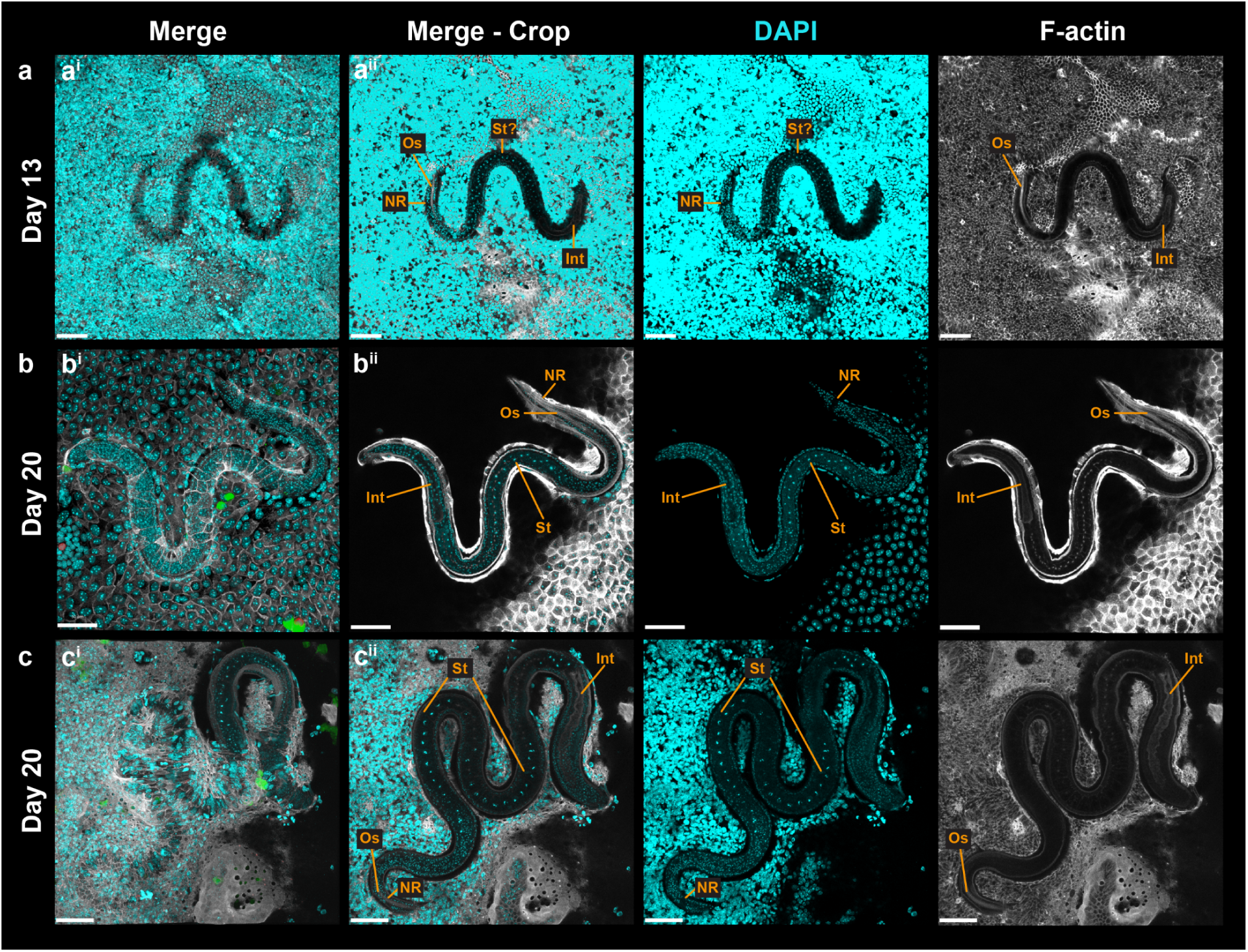
*In vitro* development of *Trichuris muris* through L2 and L3 stages. Representative confocal immunofluorescence images of caecaloids infected with *T. muris* for (a) 13 days, showing early L2 stage, and (b and c) 20 days, with parasites in a late L2/L3 stage. Complete z-stack projections (i) show intracellular worms within syncytial tunnels, and selected and cropped volumes (ii) enable visualisation of structures including NR = Nerve ring, Os = Oesophagus, St? = developing Stichosome, St = Stichosome and Int = Intestine. In aqua, DAPI stains nuclei of IECs and larvae; in green, the lectins UEA and SNA bind mucins in goblet cells; in red, Ki-67 labels dividing cells in (b) and Dckl-1 stains tuft cells in (c); and in white, phalloidin binds to F-actin. Scale bars 50 µm.

**Figure 8.**
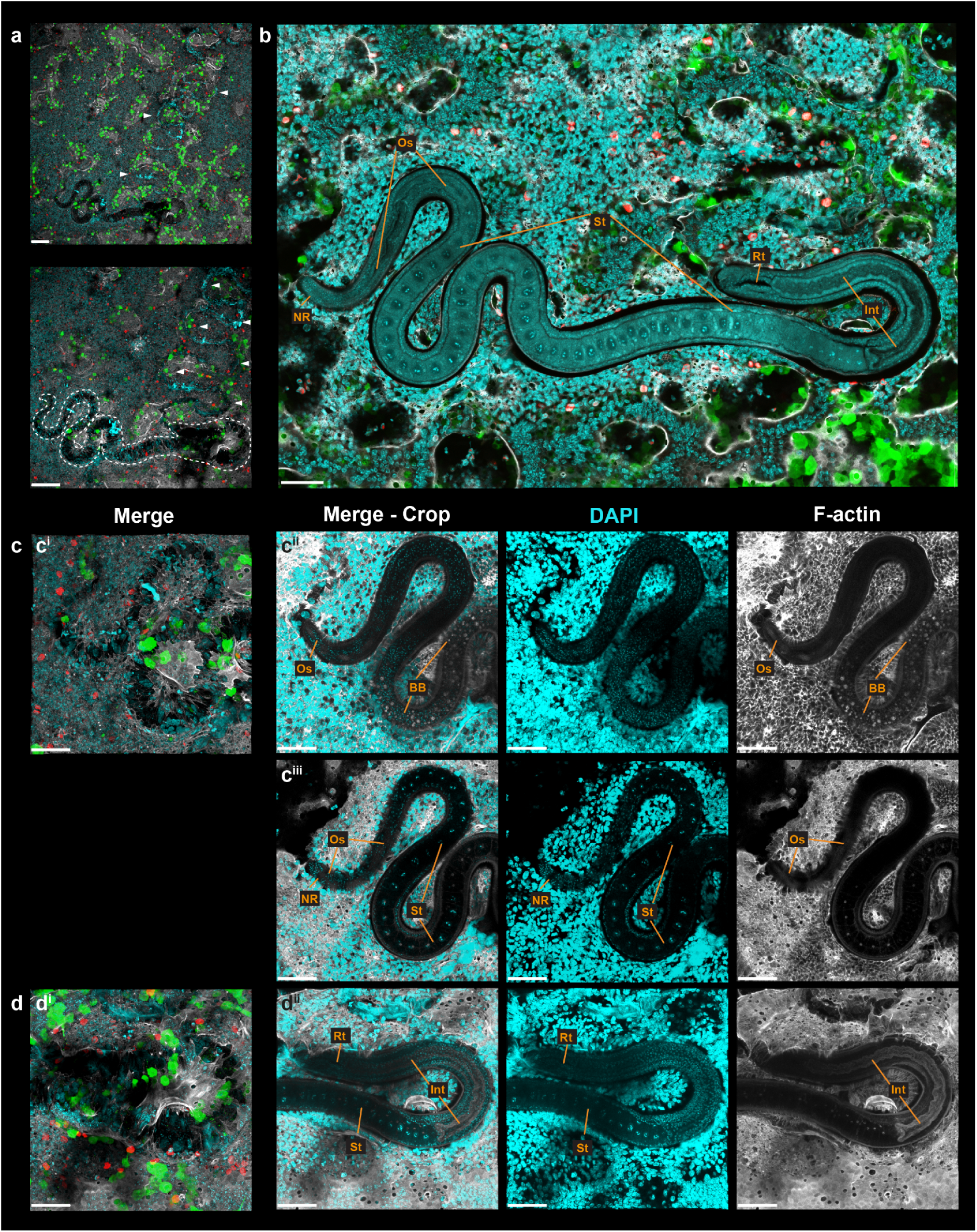
*In vitro* development of *Trichuris muris* up to L3/L4 stage. Confocal microscopy images of representative *T. muris* at L3/L4 stage from caecaloids infected for 20 days. Complete z-stack projections (a, c^i^ and d^i^) show intracellular worm within syncytial tunnel, and selected and cropped volumes (b, c^ii^, c^iii^ and d^ii^) enable visualisation of structures in the entire worm (b) and at the anterior (c) and posterior (d) end of the parasite including NR = Nerve ring, Os = Oesophagus, BB = Bacillary band, St = stichocytes forming the stichosome, Int = Intestine and Rt = Rectum. Arrowheads show syncytial tunnels. Dashed lines show larval boundaries. In aqua, DAPI stains nuclei of IECs and larvae; in green, the lectins UEA and SNA bind mucins in goblet cells; in red, Ki-67 labels dividing cells, and in white, phalloidin binds to F-actin. Scale bars in (a) 100 µm and (b-d) 50 µm.

Altogether, these results demonstrate that the caecaloid epithelium is sufficient to trigger and support sustained intracellular growth and morphological development of *T. muris* larvae *in vitro*. Parasites cultured within caecaloids follow a growth trajectory and developmental programme broadly comparable to that observed *in vivo*, acquiring increasing anatomical complexity and possessing specialised structures characteristic of later larval stages. These findings establish caecaloids as a robust platform for studying whipworm infection and development.

## Discussion

Whipworms are intracellular parasites that establish an unusual syncytial niche within the host intestinal epithelium. While other clade I nematodes such as *Trichinella spiralis* can complete key stages of their intracellular development when cultured in epithelial cell lines such as Caco-2 cells (ManWarren et al., 1997; Li et al., 1998; Gagliardo et al., 2002; Ren et al., 2011; Ming et al., 2016), whipworms appear to have more stringent requirements. Previous attempts to culture *T. muris* larvae in media or conventional epithelial cell lines have shown that newly hatched L1 larvae can survive but fail to grow or moult (White, 2016). These observations suggest that whipworms require a more complex epithelial environment than that provided by homogeneous cell lines.

In this study, we demonstrate that caecal organoids (caecaloids) provide a physiologically relevant *in vitro* environment that supports not only the invasion of intestinal epithelial cells by *T. muris* larvae but also their subsequent growth and morphogenetic development. This represents the first demonstration of the developmental progression of a parasitic nematode within an organoid system. Importantly, larvae cultured within caecaloids underwent two distinct moulting events, corresponding to the L1-L2 and L2-L3 transitions. These findings indicate that the caecaloids provides cues that are sufficient to trigger major developmental and morphogenetic events of the parasite life cycle.

The ability of caecaloids to support parasite development likely reflects their capacity to recapitulate the heterogeneity in cellular populations and spatial organisation of the intestinal epithelium. Unlike epithelial cell lines, intestinal organoids contain multiple differentiated epithelial cell types, including enterocytes, goblet cells, enteroendocrine cells and tuft cells, organised in a crypt-like architecture (Duque-Correa et al., 2020b). This cellular and spatial heterogeneity may provide the mechanical and biochemical signals required for parasite invasion and development. The observation that larvae not only increase in size but also develop complex morphological structures, including specialised features such as the stichosome and the bacillary band, suggests that these host-derived cues are sufficient to support key aspects of whipworm morphogenesis.

To robustly validate our caecaloid system as a platform to faithfully study *T. muris* infection and development, we performed a thorough biometrical analysis comparing parasite growth and development *in vivo* and *in vitro*. Much of the existing biometrical data describing the growth and developmental progression of *T. muris* originates from studies published several decades ago (Fahmy, 1954; Wakelin, 1969; Panesar, 1989; Blasco-Costa and Poulin, 2017; Panti-May et al., 2023), with the risks of data being out of date due to potential genetic drift in parasite laboratory strains and host mouse strain-dependent developmental variation. Therefore, in this study, we generated a comprehensive and up-to-date dataset of the gross growth and morphological development of *T. muris in vivo*, which served as a reference framework against which parasite growth and development within caecaloids were directly evaluated. We found that *in vitro* parasite development in caecaloids remains less robust than *in vivo*. Although larvae grew and underwent moulting within caecaloids, overall developmental progression was reduced compared with infections in the host intestine. This observation suggests that there are additional cues in the *in vivo* environment that are absent or incompletely recapitulated in the caecaloid system. Such cues may include biomechanical forces, interactions with the microbiota, or signals derived from non-epithelial cell types such as immune or mesenchymal cells.

To identify host factors that regulate parasite development, we will next perform side-by-side transcriptomic profiling of larvae developing in the host caecum and within caecaloids. This will reveal parasite pathways that are differentially activated between the two conditions. Such analyses will help pinpoint specific host-derived signals that promote moulting and growth, guiding the refinement of organoid culture systems to ultimately allow the full life cycle of whipworms to be recapitulated *in vitro*.

A recent study by Jung et al. (2025) reported preliminary evidence for *T. muris* growth *in vitro* within colon organoids (colonoids). However, the analysis was restricted to only 14 days post infection and primarily focused on increases in larval size. In contrast, our findings provide clear definitive evidence that larvae cultured within caecaloids undergo both sustained growth and morphological development. We demonstrate the development of highly specialised structures such as the stichosome and bacillary band and document parasite development up to day 20 post-infection where larvae reached a maximum length of over 1600 μm and a high level of physiological complexity. This likely reflects the fact that the caecum is the natural site of infection for *T. muris*, and that organoids derived from this tissue may therefore more accurately reproduce the *in vivo* epithelial niche of the parasite.

### Outlook - application

Together, our findings establish caecaloids as a powerful experimental platform for studying whipworm biology. By providing a physiologically relevant *in vitro* environment that supports both epithelial invasion and developmental progression, this system opens new opportunities to dissect the molecular and cellular mechanisms governing host–parasite interactions in the intestinal epithelium.

Despite its current limitations, the caecaloid infection model provides several advantages for studying host–parasite interactions. Firstly, the model offers the potential to reduce reliance on animal experiments while enabling controlled manipulation of both host and parasite components in a reductionistic system. Caecaloids enable the direct interactions of whipworms with the IE to be dissected and uncoupled from the changes to this tissue when the immune system responds to infection. By introducing other cellular populations present in the intestine or soluble mediators experimentally, their impact on whipworm development can be evaluated in a controlled manner.

Another advantage of the caecaloid model is that the entire parasite can be visualised in contact with its host cells, which is nearly impossible for *in vivo* infections where whipworms often infect several crypts that cannot be captured in a single section. Spatial transcriptomics of whipworm-infected caecaloids will enable simultaneous analysis of host and parasite, revealing the precise cellular context of gene expression changes *in situ* and, thus, a comprehensive investigation of the interactions of host IECs with the entire body of the parasite. These data will also reveal the tissue organisation, structure, and potential function of whipworm organs that control parasite feeding, sensory and secretory behaviours, shedding light on how the parasites live inside their host.

Furthermore, the *in vitro* system allows parasite development to be monitored at a much higher temporal resolution than it is possible in animal models, enabling larvae to be recovered at more defined time points and developmental stages. Of note, the slower development of *T. muris* within the caecaloid infection model unorthodoxically offers a unique opportunity to observe the growth of the worm and its tissues in “slow motion”, allowing for unprecedented temporal resolution of these processes. This will facilitate detailed investigation of early developmental transitions and the cellular mechanisms underlying epithelial invasion.

## Data and code availability

Raw data such as image sets and measurements are available upon request.

## Acknowledgements - Funding

This work was supported by the Sir Henry Dale Fellowship jointly funded by the Wellcome Trust and the Royal Society (222546/Z/21/Z, M.A.D-C.); the Isaac Newton Trust (22.39(f), M.A.D-C.), the Wellcome Trust (206194, M.B; 203151/Z/16/Z, M.A.D-C) and the UKRI Medical Research Council (MC_PC_17230, M.A.D-C). SRD is supported by a UKRI Future Leaders Fellowship (MR/T020733/1). The funders had no role in study design, data collection and analysis, decision to publish, or preparation of the manuscript. For the purpose of Open Access, the author has applied a CC BY public copyright licence to any Author Accepted Manuscript version arising from this submission.

## Supplementary materials

**Supplementary Figure 1.**
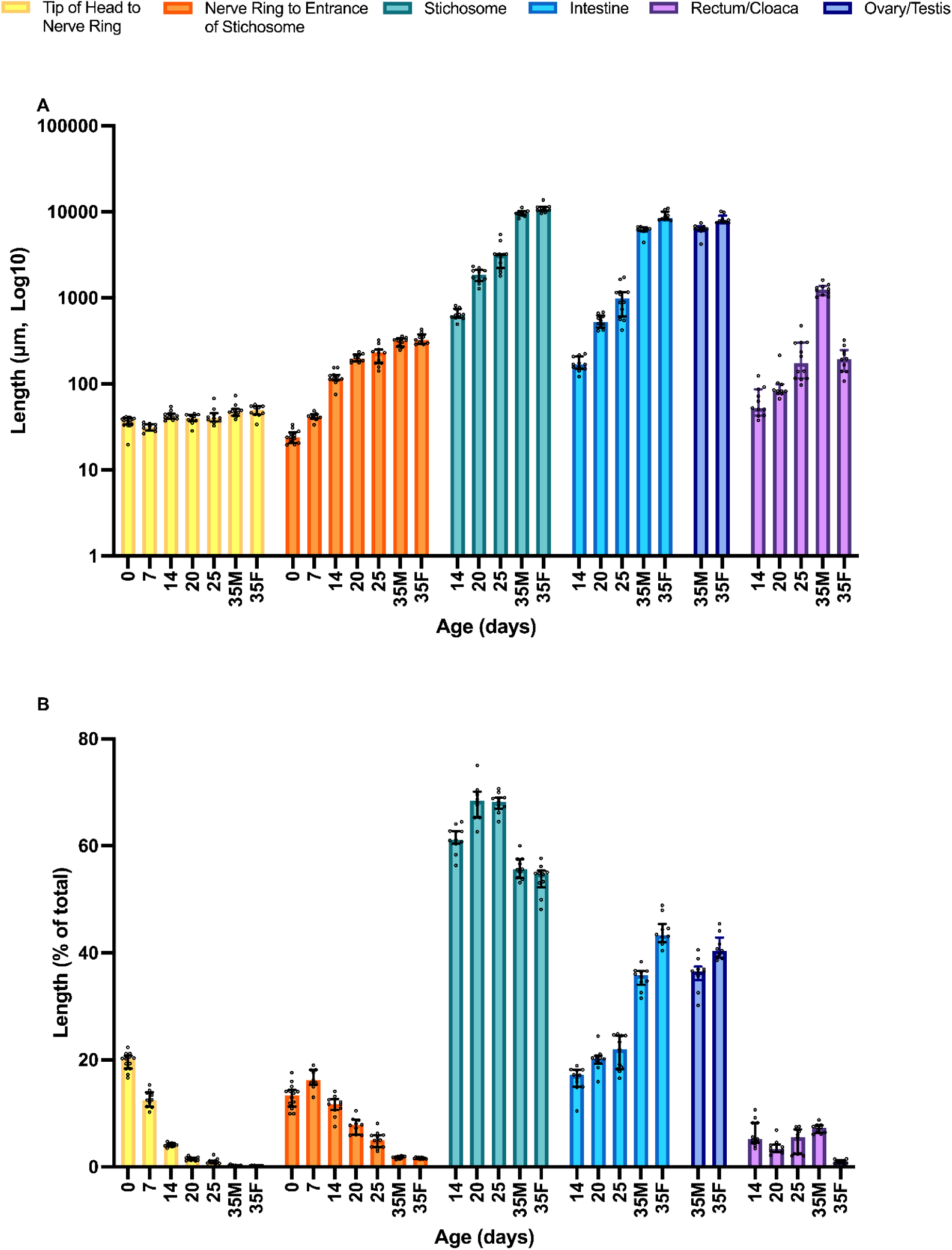
Absolute and relative measurements of *Trichuris muris* substructures across development. (A) Length of tip of head to nerve ring (NR), length from NR to entrance of stichosome (or to end of oesophagus for day 0 and day 7), stichosome (ST), intestine (IN), ovary/testis (OV/TES), and rectum/cloaca (REC/CL) were measured on *T. muris* at days 0 (n_NR_ = 17, n_NR to ST_ = 14), 7 (n_NR_ = 10, n_NR to ST_ = 9), 14 (n_NR_ = 11, n_NR to ST_ = 11, n_ST_ = 11, n_IN_ = 11, n_REC_ = 11), 20 (n_NR_ = 10, n_NR to ST_ = 10, n_ST_ = 10, n_IN_ = 10, n_REC_ = 10), 25 (n_NR_ = 10, n_NR to ST_ = 11, n_ST_ = 12, n_IN_ = 12, n_REC_ = 12), and 35 (females (n_NR_ = 10, n_NR to ST_ = 9, n_ST_ = 10, n_IN_ = 10, n_REC_ = 10, n_OV_ = 9) and males (n_NR_ = 10, n_NR to ST_ = 10, n_ST_ = 10, n_IN_ = 10, n_CL_ = 10, n_TES_ = 10) p.i.. Median with individual data points and interquartile range are shown. (B) Tissue lengths expressed as a percentage of total body length from *T. muris* recovered at days 0 (n_NR_ = 14, n_NR to ST_ = 14), 7 (n_NR_ = 10, n_NR to ST_ = 9), 14 (n_NR_ = 11, n_NR to ST_ = 11, n_ST_ = 11, n_IN_ = 11, n_REC_ = 11), 20 (n_NR_ = 10, n_NR to ST_ = 10, n_ST_ = 10, n_IN_ = 10, n_REC_ = 10), 25 (n_NR_ = 10, n_NR to ST_ = 11, n_ST_ = 11, n_IN_ = 11, n_REC_ = 11),, and 35 (females (n_NR_ = 10, n_NR to ST_ = 9, n_ST_ = 10, n_IN_ = 10, n_REC_ = 10, n_OV_ = 9) and males (n_NR_ = 10, n_NR to ST_ = 10, n_ST_ = 10, n_IN_ = 10, n_CL_ = 10, n_TES_ = 10) p.i.. Median with individual data points and interquartile range are shown.

## Notes

### Competing Interest Statement

The authors have declared no competing interest.

